# Pre-existing SIV infection increases expression of T cell markers associated with activation during early *Mycobacterium tuberculosis* co-infection and impairs TNF responses in granulomas

**DOI:** 10.1101/2020.12.14.422668

**Authors:** Erica C. Larson, Amy L. Ellis, Mark A. Rodgers, Alexis J. Balgeman, Ryan V. Moriarty, Cassaundra Ameel, Tonilynn Baranowski, Jaime Tomko, Chelsea Causgrove, Pauline Maiello, Shelby L. O’Connor, Charles A. Scanga

**Affiliations:** Department of Microbiology and Molecular Genetics and Center for Vaccine Research, University of Pittsburgh School of Medicine, Pittsburgh, Pennsylvania, United States of America; Center for Vaccine Research, University of Pittsburgh School of Medicine, Pittsburgh, Pennsylvania, United States of America; Department of Pathology and Laboratory Medicine, University of Wisconsin - Madison, Wisconsin, United States of America; Wisconsin National Primate Research Center, University of Wisconsin - Madison, Wisconsin, United States of America

## Abstract

Tuberculosis (TB) is the leading infectious cause of death among people living with HIV (PLHIV). PLHIV are more susceptible to contracting *Mycobacterium tuberculosis* (*Mtb*) infection and often have worsened TB disease. Understanding the immunologic defects caused by HIV and the consequences it has on *Mtb* co-infection is critical in combating this global health epidemic. We previously established a model of simian immunodeficiency virus (SIV) and *Mtb* co-infection in Mauritian cynomolgus macaques (MCM), and showed that SIV/*Mtb* co-infected MCM had rapidly progressive TB. We hypothesized that pre-existing SIV infection impairs early T cell responses to *Mtb* infection. To test our hypothesis, we infected MCM with SIVmac239 intrarectally followed by co-infection with a low dose of *Mtb* Erdman 6 months later. SIV-naïve MCM were infected with *Mtb* alone as controls. Six weeks after *Mtb* infection, animals were necropsied and immune responses were measured by multiparameter flow cytometry. While the two groups exhibited similar TB progression at time of necropsy (Nx), longitudinal sampling of the blood (PBMC) and airways (BAL) revealed a significant reduction in circulating CD4+ T cells and an influx of CD8+ T cells in airways following *Mtb* co-infection of SIV+ animals. Differences in the activation markers CD69, PD-1, and TIGIT were observed. At sites of *Mtb* infection (*i.e.* granulomas), SIV/*Mtb* co-infected animals had a higher proportion of CD4+ and CD8+ T cells expressing PD-1 and TIGIT. In addition, there were fewer TNF-producing CD4+ and CD8+ T cells in granulomas and airways of SIV/*Mtb* co-infected animals. Taken together, we show that concurrent SIV infection alters T cell phenotypes in granulomas during the early stages of TB disease. As it is critical to establish control of *Mtb* replication soon after infection, these phenotypic changes may distinguish the immune dysfunction that arises from pre-existing SIV infection which promotes TB progression.

**Author Summary:** People living with HIV are incredibly susceptible to TB and, when co-infected with *Mtb*, often develop serious TB disease. We do not yet understand precisely how HIV infection impairs the early stages of the adaptive immune response against *Mtb* bacilli. We employed a non-human primate model of HIV, using SIV as a surrogate for HIV, followed by *Mtb* co-infection to investigate the immunologic defects associated with pre-existing SIV infection over the first six weeks of *Mtb* co-infection. Our study focused on CD4+ and CD8+ T cells as these cells are known to play an important role in *Mtb* control. We found more CD8+ T cells in granulomas, the sites of *Mtb* infection, from SIV/*Mtb* co-infected animals, with little difference in CD4+ T cells. SIV/*Mtb* co-infected animals and animals infected with SIV alone had a higher proportion of both CD4+ and CD8+ T cells expressing activation markers compared to SIV-naïve animals, consistent with SIV-dependent immune activation. Notably, we observed a lower proportion of TNF-producing T cells, a cytokine critical for *Mtb* control, in granulomas and airways of SIV/*Mtb* co-infected animals. Taken together, these data show that pre-existing SIV alters T cell phenotypes and reduces TNF responses early in *Mtb* infection.

## Introduction

Tuberculosis (TB) is a major global health concern, especially among people living with HIV (PLHIV). TB, caused by the bacterium *Mycobacterium tuberculosis* (*Mtb*), is the leading cause of death worldwide among PLHIV, accounting for one-third of AIDS deaths [1]. PLHIV are incredibly susceptible to *Mtb* and have a 20-fold greater risk of developing TB than HIV-naïve individuals [1]. PLHIV have a greater risk of both developing active TB disease or reactivating a latent TB infection [2, 3]. The risk for contracting bacterial infections, including *Mtb*, is higher in PLHIV even before circulating CD4+ T cell counts fall [4–7], although susceptibility rapidly increases as CD4+ T cell counts decline [8]. Furthermore, the TB risk remains elevated even after individual are put on antiretroviral therapy [9]. Thus, there is a critical need to identify more precisely the underlying mechanisms by which pre-existing HIV infection impairs the immune response to *Mtb*.

Knowledge gaps remain in our understanding of early immune events following *Mtb* infection [10]. T cells are an important component of the adaptive immune response and play a critical role in controlling *Mtb* [11, 12]. In both humans [13] and non-human primates (NHP) [14], T cell responses to *Mtb* take several weeks to fully develop [13]. However, early responses to *Mtb* and how they are affected by HIV is difficult to study in humans as the exact time points of both HIV and/or *Mtb* infection are often unknown. NHP are an ideal model in which to study interactions between HIV and TB. NHP are susceptible to SIV infection, a simian surrogate for HIV, and they closely recapitulate key features of TB disease in humans [15–20]. They offer wider access to tissues often inaccessible in human studies and provide insight into localized immunologic responses that are not necessarily reflected in blood [21–24].

Several NHP studies of the effect of SIV infection on reactivation of latent TB infection showed that SIV decreased CD4+ T cells in lungs, dysregulated cytokine responses, increased *Mtb* dissemination and elevated rates of TB reactivation [25–31]. However, fewer studies have investigated the effects of pre-existing SIV infection on the outcome of *Mtb* co-infection. In animals with an acute SIV infection (6 weeks) followed by *Mtb* co-infection, TB disease was more severe and SIV+ animals succumbed to their disease much earlier than their counterparts infected with *Mtb* only [32]. Early mortality in SIV-infected animals also has been reported following *M. bovis* co-infection [33, 34]. SIV infection, whether in the context of pre-existing latent TB infection or prior to *Mtb* co-infection, impairs the immune system and leads to more extensive and severe TB disease. However, the precise nature of this immunologic impairment, as well as the mechanisms by which it occurs, have yet to be fully elucidated.

We recently established a model of SIV and *Mtb* co-infection in Mauritian cynomolgus macaques (MCM) [35]. We showed that animals with a pre-existing SIV infection exhibited rapid TB progression upon co-infection with *Mtb* and reached clinical endpoint considerably sooner (9 - 13 weeks post-*Mtb* co-infection) than SIV-naïve animals. Strikingly, the number of pulmonary granulomas detected by PET/CT imaging increased significantly between 4 and 8 weeks following *Mtb* co-infection in SIV+ animals, while the numbers remained more stable SIV-naïve animals [35]. This timeframe also corresponds to the period when the T cell response to *Mtb* infection of NHP develops fully [14]. We thus hypothesized that the period between 4 and 8 weeks after *Mtb* co-infection may be the critical window during which immunologic defects associated with pre-existing SIV infection manifest to alter the trajectory of the TB disease.

Here, we characterized differences in conventional CD4+ and CD8+ T cell responses at 6 weeks after *Mtb* infection in SIV-naïve and SIV+ MCM, at a time that corresponds to when the trajectory of TB disease began to diverge in SIV-naïve and SIV+ MCM [35]. By 6 weeks post-infection, the adaptive immune response against *Mtb* has developed [14, 19] and granulomas are just beginning to acquire mycobacterial killing capacity [36]. This time point also precedes the appearance of extensive necrosis and other immunopathologies associated with advanced TB that would confound immunologic analysis of T cell responses. We investigated changes in conventional T cells across several compartments: the airways, where *Mtb* is first encountered; blood, a reflection of cell trafficking and circulating immune cells that is easily sampled; and lung tissue including individual granulomas, as local sites of *Mtb* infection. We observed a decrease in the CD4:CD8 ratio across all these compartments in SIV+ animals. SIV infection led to T cell activation in mucosal tissue as indicated by increased frequencies of PD-1 and TIGIT expression on CD4+ T cells and, to a lesser extent, on CD8+ T cells, in lung tissue and granulomas. Lastly, there were fewer TNF-producing CD4+ T cells and CD8+ T cells in granulomas from SIV/*Mtb* co-infected animals compared to animals infected with *Mtb* alone. Taken together, these data provide novel insights into the mechanism by which SIV dysregulates the adaptive immune response to *Mtb* infection that may promote TB progression.

## Results

### No difference in TB disease pathology or bacterial burden in SIV+ and SIV-naïve animals at 6 weeks post-*Mtb* infection

We previously observed rapid progression of TB disease as measured by 2-deoxy-2-(18F) fluorodeoxyglucose (FDG) PET/CT imaging beginning 4 - 8 weeks after *Mtb* co-infection of SIV+ MCM [35]. This suggests that this period is critical in determining the trajectory of TB disease- either bacterial containment and TB control or bacterial dissemination and progressive TB. In the current study, we investigated the immunologic defect caused by SIV that increases *Mtb* susceptibility by comparing two experimental groups: MCM infected with *Mtb* only and SIV+ MCM co-infected with *Mtb*. Additionally, data from a small subset of MCM (n=4) infected only with SIV was included as a comparator for some analyses (Fig 1A). Necropsies were conducted 6 weeks after *Mtb* infection and, as expected, neither overall TB pathology nor total thoracic bacterial burden differed between the two groups (Fig 1B, C). Nor were differences observed in TB pathology and bacterial burden in individual compartments, including lung, thoracic lymph nodes, and extrapulmonary organs (S1 Fig A-E). There also was no observable difference in the frequency of *Mtb* culture-positive granulomas (S1 Fig F), indicating that granulomas had not yet developed substantial *Mtb* killing capacity at 6 weeks post-*Mtb* infection, regardless of the animals’ SIV status. This is not surprising as the total bacterial burden in Chinese cynomolgus macaques peaks between 4 and 6 weeks post-*Mtb* and is then subsequently reduced as mycobactericidal activity develops [36, 37]. Taken together, these data suggest that TB disease state in MCM with chronic SIV infection resembles that in SIV-naïve animals at 6 weeks post-*Mtb* infection, even though TB disease trajectory soon diverges in SIV+ animals [35]. Thus, 6 weeks post-*Mtb* represents an ideal time point to compare the conventional T cell response between SIV+ and SIV-naïve MCM - a time when adaptive immunity to *Mtb* appears but prior to the development of extensive immunopathology in SIV+ animals which confounds analysis [35].

**Fig 1.**
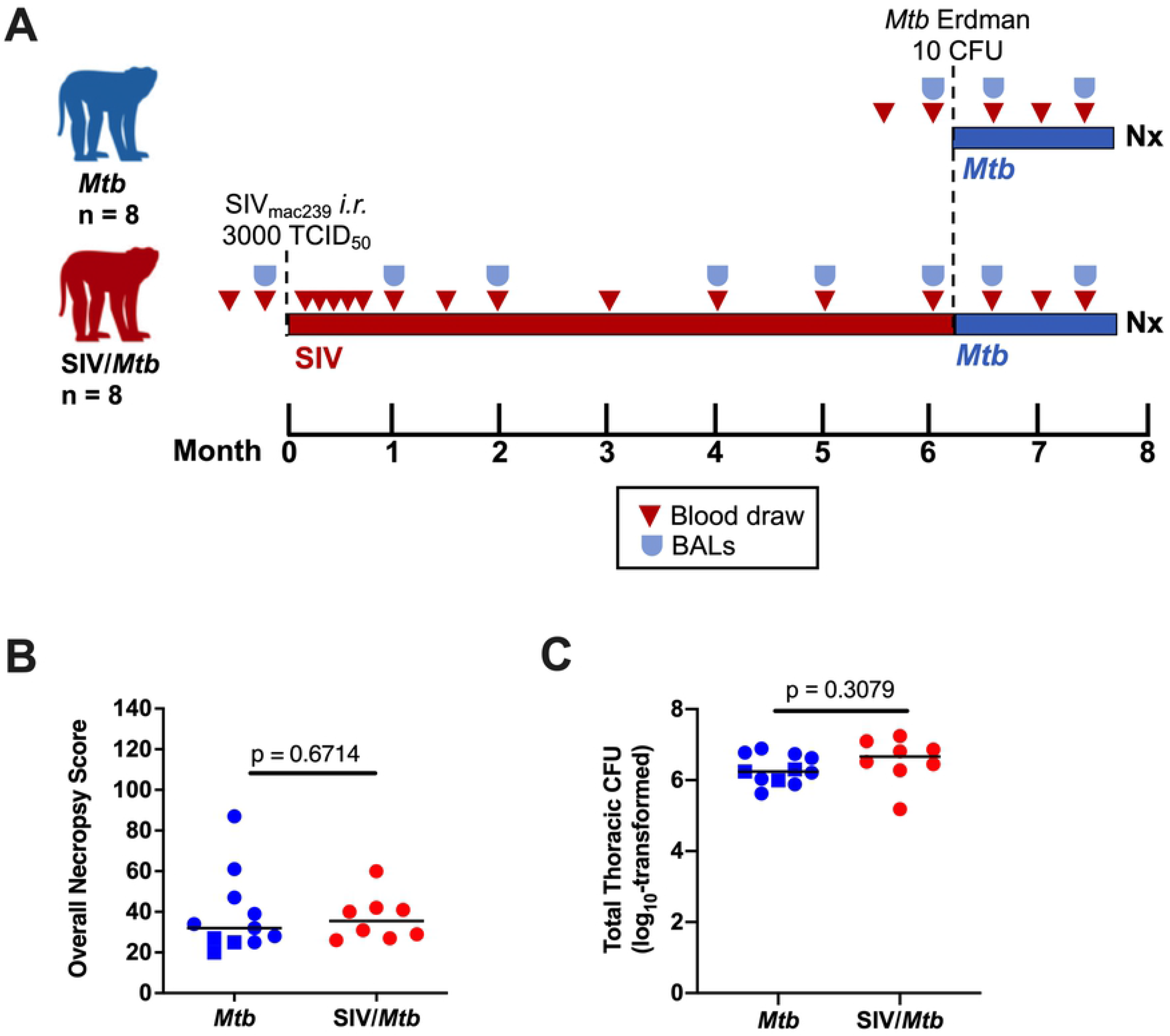
No difference in overall Necropsy (Nx) score and total thoracic CFU. A.) Experimental design. Three cohorts of animals: *Mtb* only (blue, n = 11); SIV/*Mtb* (red, n = 8); and SIV only (black, n = 4). B.) Nx samples from *Mtb*-infected animals were scored and compiled to quantify TB pathology. Medians are shown. *Mtb* only (blue) and SIV/*Mtb* (red). Mann-Whitney tests were used to determine statistical significance. C.) Total Thoracic CFU combined CFU from lung tissue, thoracic lymph nodes, and granulomas from *Mtb*-infected animals. Medians are shown. An unpaired t test was used to determine statistical significance.

### Pre-existing SIV infection decreases the CD4:CD8 T cell ratio across multiple tissue compartments

A hallmark of HIV infection is the loss of CD4+ T cells, resulting in a decreased CD4:CD8 T cell ratio [38, 39]. The CD4:CD8 ratio of circulating T cells in SIV+ animals was significantly decreased, both pre- and post-*Mtb* co-infection (Fig 2A). Similarly, the CD4:CD8 T cell ratio in airways of SIV+ animals was significantly lower over the course of *Mtb* co-infection than compared to SIV-naïve animals (Fig 2B). In tissue samples harvested at necropsy, CD4:CD8 ratios were significantly lower in randomly collected lung tissue and in individual granulomas (Fig 2C, D). The CD4:CD8 T cell ratio in lung tissue from the 4 MCM infected with SIV alone was similar to that observed in the SIV/*Mtb* animals (Fig 2C), suggesting that the altered ratio was a consequence of SIV, rather than the *Mtb* co-infection. Thus, the CD4:CD8 ratio was lower in SIV+ animals across all tissue compartments measured.

**Fig 2.**
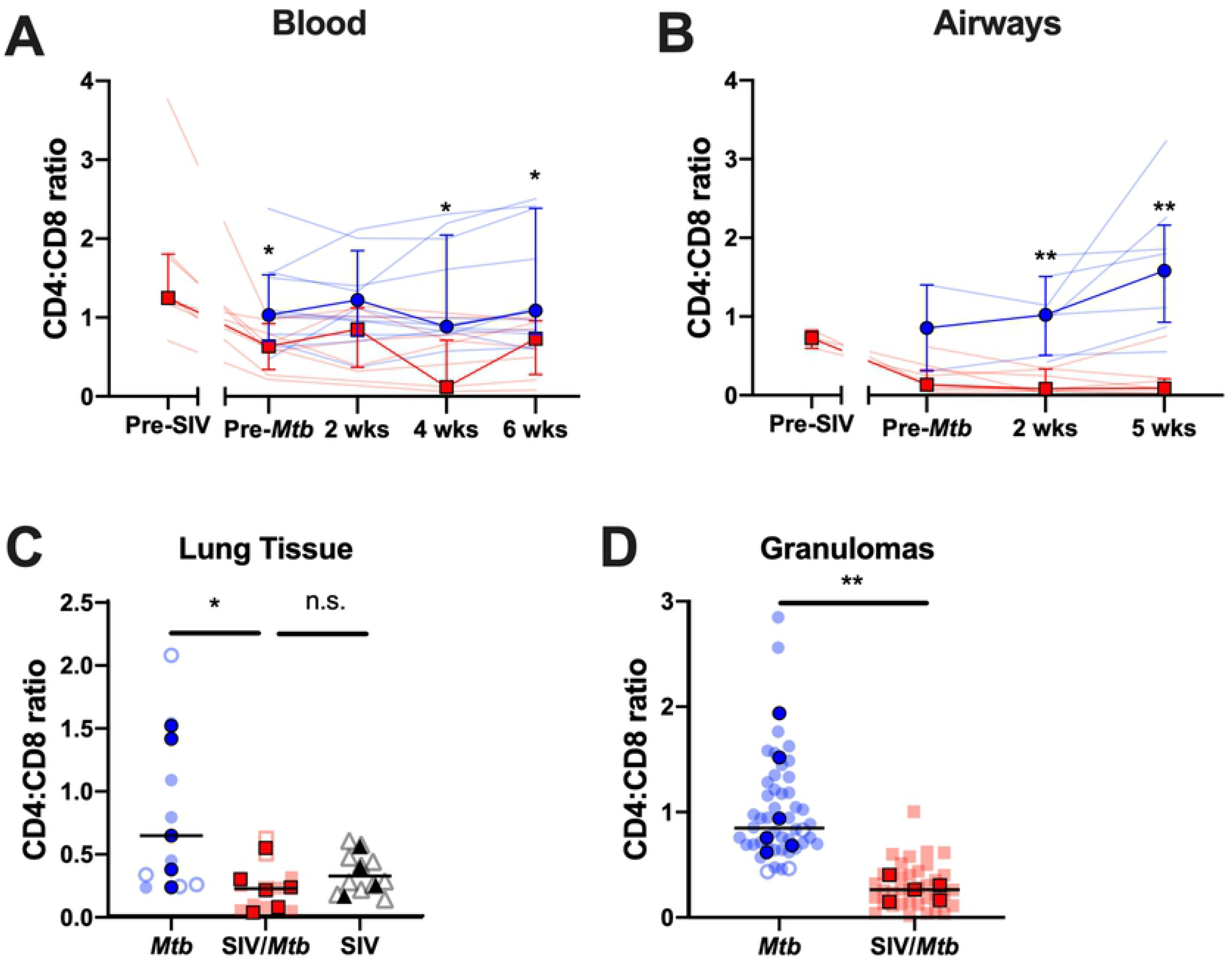
Pre-existing SIV systemically decreases CD4:CD8 ratios. Blood (A.) and airway (B.) samples were collected over the course of the study and cells were stained for flow cytometry. Data from *Mtb*-only animals is shown in blue; and SIV/*Mtb* co-infected animals are in red. Solid lines indicate median with interquartile range. Lighter lines indicate CD4:CD8 ratios of individual animals. Mann-Whitney tests were performed at each time point between SIV+ and SIV-naïve groups to determine statistical significance. Lung tissue (C.) and granulomas (D.) were collected at necropsy and stained for flow cytometry. *Mtb*-only animals are in blue, SIV/*Mtb* co-infected animals in red, and SIV-only animals in black. Darker symbols represent the median of each animal and the lighter symbols are individual samples. Closed symbols represent CFU+ tissue and open symbols are sterile (CFU-) tissue. For lung tissue, a Kruskal-Wallis test was performed with a Dunn’s multiple comparison follow-up test comparing mean ranks to SIV/*Mtb*. For granulomas, a Mann-Whitney test was performed to determine statistical significance. Statistical significance indicated: * p < 0.05; ** p < 0.01.

### SIV alters the number of T cells in circulation, in the airways, and in granulomas

Since a lower CD4:CD8 ratio itself does not differentiate whether the change is due to decreased CD4+ T cells or increased CD8+ T cells, we determined the absolute numbers of these two cell types in different tissue compartments. Over the course of HIV infection in humans, circulating CD4+ T cell counts can decline to ≤200 CD4+ T cells/μL blood, an AIDS-defining criterion [40]. In our MCM model, the SIV+ macaques had fewer circulating CD3+ T cells than SIV-naïve animals, both before and after *Mtb* co-infection (Fig 3A). This decrease was largely due to significantly fewer circulating CD4+ T cells in the SIV+ animals (Fig 3B). CD8+ T cell counts in the blood were also lower, but not statistically significantly (Fig 3C).

**Fig 3.**
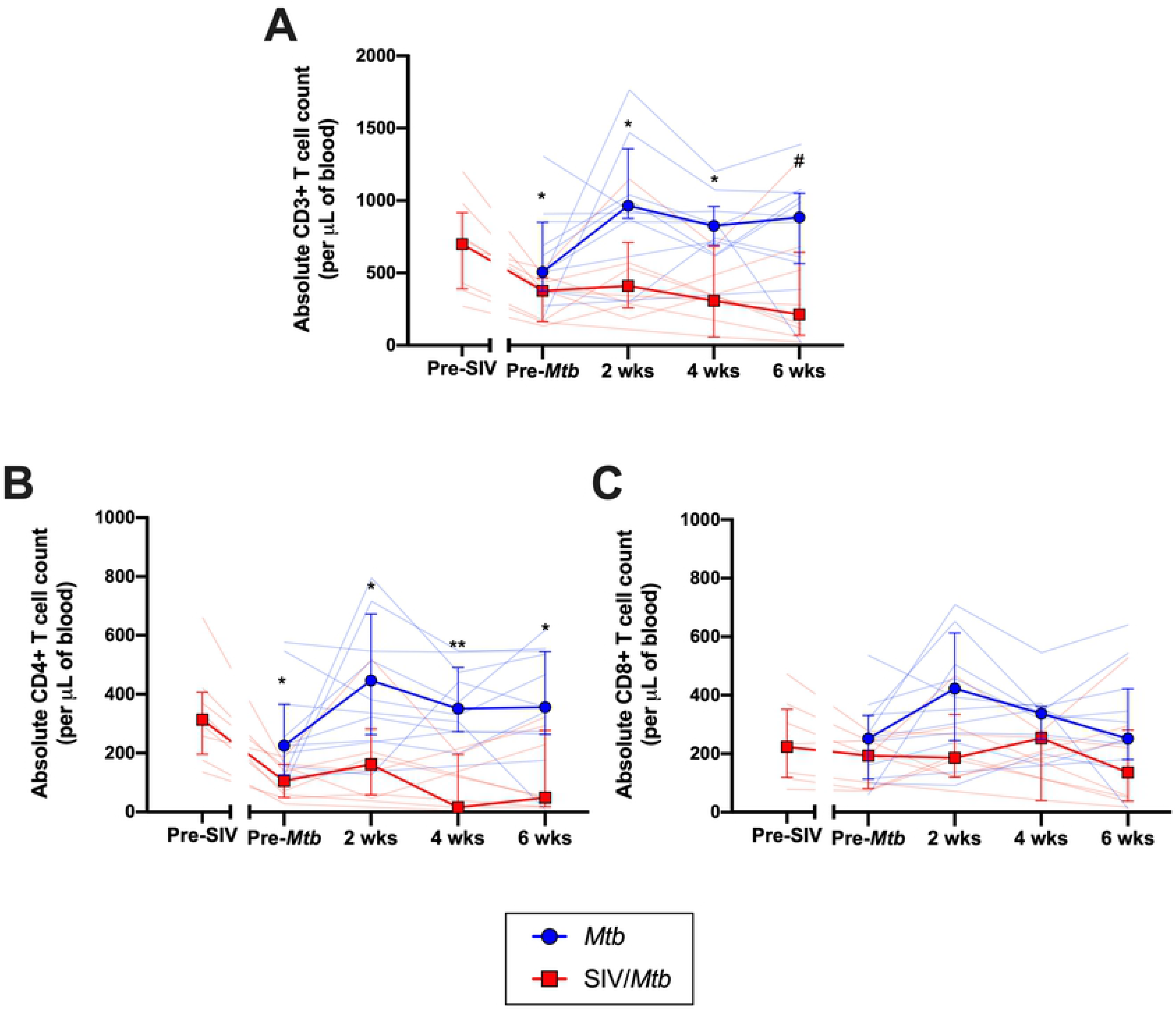
SIV decreases peripheral T cell populations prior to *Mtb* challenge and impairs their expansion following *Mtb* challenge. PBMC were collected throughout the course of SIV and *Mtb* infection. *Mtb*-only animals are in blue; SIV/*Mtb* co-infected animals in red. Solid lines indicate median with interquartile range. Lighter lines indicate absolute cell counts of individual animals. Mann-Whitney tests were performed to determine statistical significance between SIV+ and SIV-naïve groups at indicated time points. A.) Absolute CD3+ T cells counts in PBMC. B.) Absolute CD4+ T cell counts in PBMC. C.) Absolute CD8+ T cell counts in PBMC. Statistical significance indicated: # 0.05 < p < 0.1; * p < 0.05; ** p < 0.01.

There were slightly more total CD3+ T cells in the airways of SIV+ animals following *Mtb* co-infection than in airways of animals infected with *Mtb* only (~0.4 log_10_ difference at 2 and 5 weeks; Fig 4A). Interestingly, the significant decrease in the airway CD4:CD8 ratio observed in SIV+ animals (Fig 2B), was not due to a loss of CD4+ T cells (Fig 4B) but rather a significant increase in CD8+ T cells (~0.7-0.8 log_10_ difference at 2 and 5 weeks post-*Mtb* infection, respectively; Fig 4C). An influx of SIV-specific CD8+ T cells into airways has been reported in acute SIV infection [41]. Although we did not measure SIV-specificity here, it is likely that the CD8+ T cells accumulating in the airways of SIV+ MCM are SIV-specific. The number of T cells in the lung tissue did not differ between SIV/*Mtb* co-infected animals and those infected with *Mtb* alone (Fig 5A). However, granulomas from SIV/*Mtb* animals showed an increase in the number of CD8+ T cells present at 6 weeks post-*Mtb* infection when compared to SIV-naïve animals, although this observation was a trend (p = 0.0552; Fig 5B).

**Fig 4.**
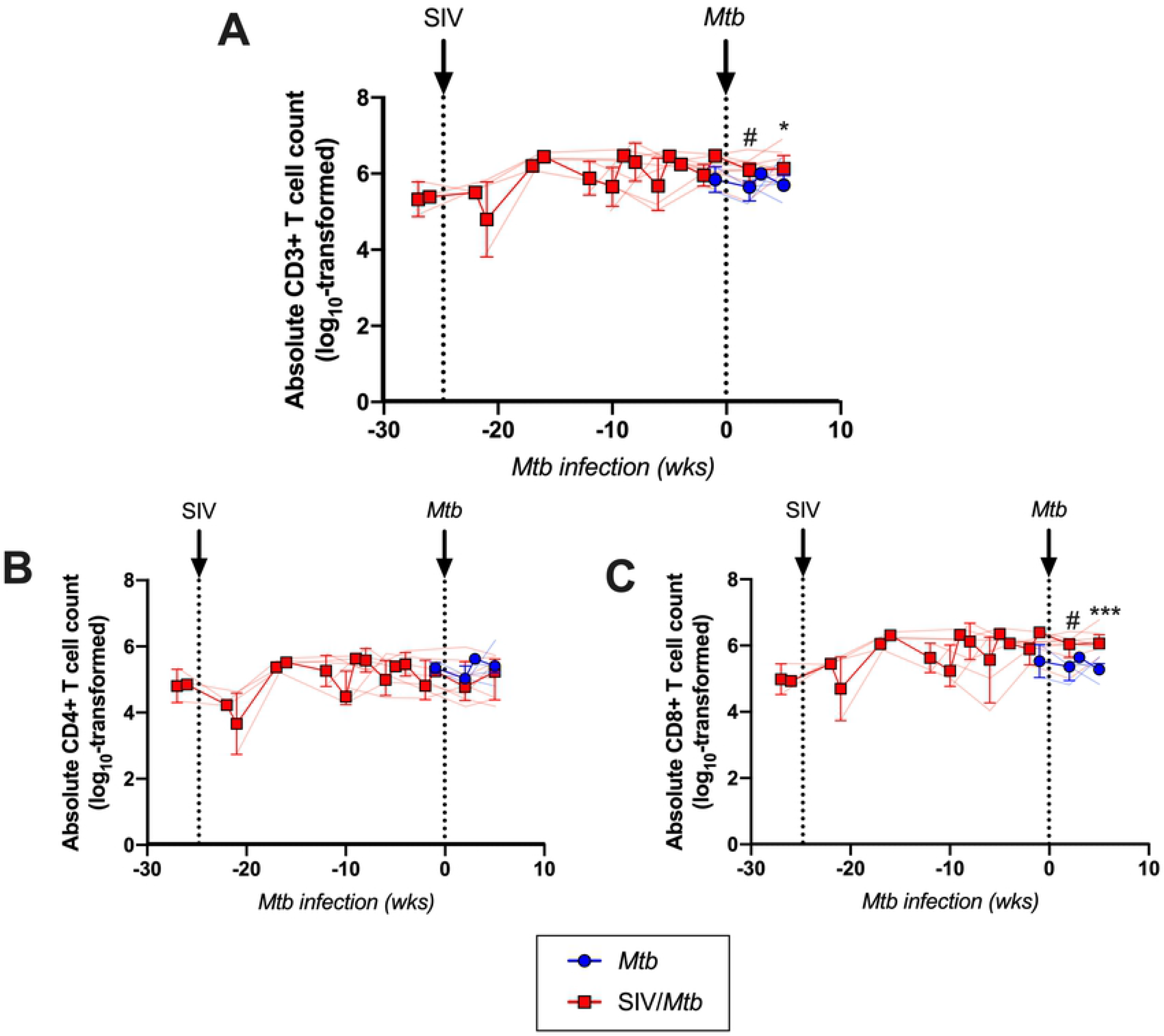
SIV increases CD8+ T cells in the airways following *Mtb* challenge. Cells from airways were collected by BAL throughout the course of SIV and *Mtb* infection. *Mtb*-only animals (blue); SIV/*Mtb* co-infected animals (red). Solid lines indicate median with interquartile range. Lighter lines indicate absolute cell counts of individual animals. A.) Absolute CD3+ T cells counts in BALs. B.) Absolute CD4+ T cell counts in BALs. C.) Absolute CD8+ T cell counts in BALs. Combined 2- and 3-week time points for the *Mtb*-only animals were used for statistical analysis. Mann-Whitney tests were performed at indicated time points between SIV+ and SIV-naïve groups to determine significance; # 0.05 < p < 0.1, * p < 0.05, & *** p < 0.001.

**Fig 5.**
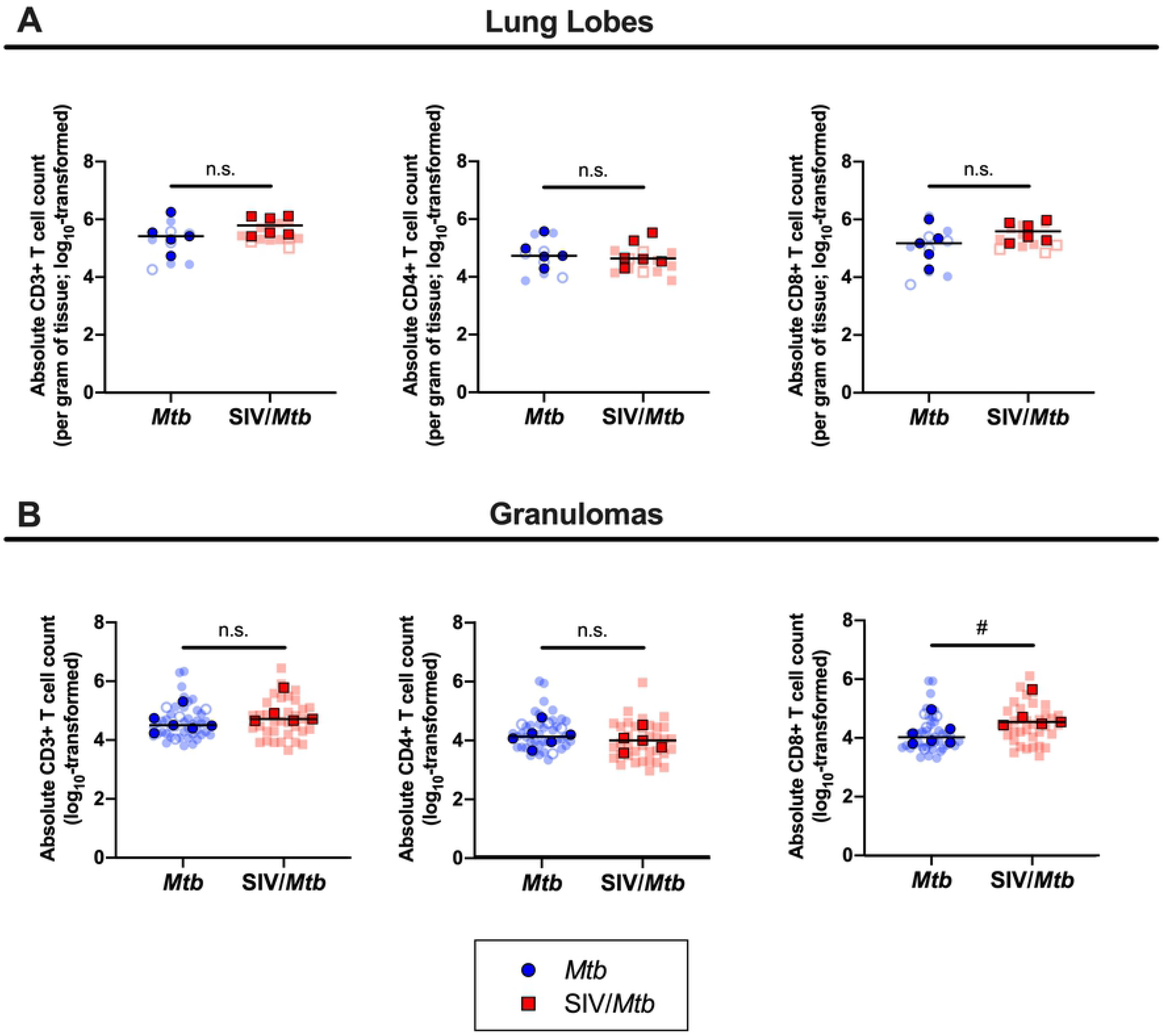
No significant differences in absolute T cells counts in lung tissue of SIV/*Mtb* animals, however CD8+ T cells are increased in granulomas. Lung tissue and granulomas were collected at necropsy and stained for flow cytometry. *Mtb*-only animals are in blue; SIV/*Mtb* co-infected animals are in red. Darker symbols represent the median of each animal and the lighter symbols are individual samples. Closed symbols represent CFU+ tissue and open symbols are sterile (CFU-). A.) Absolute T cells counts per gram of lung tissue. B.) Absolute T cell counts in granulomas. Mann-Whitney tests were performed to determine statistical significance between SIV+ and SIV-naïve groups; # 0.05 < p < 0.1.

### Circulating CD4+ and CD8+ T cells are more activated in SIV+ animals post-*Mtb* infection

Having compared CD4+ and CD8+ T cell frequencies in various compartments following *Mtb* infection of SIV+ and SIV-naïve MCM, we then assessed phenotypic differences of these cells circulating in blood from these two groups of animals. We measured several T cell activation markers including CD69, PD-1, CTLA-4, and TIGIT. The frequencies of CD4+ T cells expressing PD-1 or CD69 were similar between SIV+ and SIV-naïve animals prior to *Mtb* infection but significantly increased in the SIV+ animals during the 6 weeks following *Mtb* infection (Fig 6A, B). In contrast, the frequencies of CD8+ T cells expressing PD-1 or CD69 were higher in SIV+ animals even prior to *Mtb* infection and remained elevated during *Mtb* co-infection (Fig 6C, D). These data suggest that circulating CD4+ T cells express surface markers consistent with activation following *Mtb* infection, while the expression of activation markers on CD8+ T cells is likely a consequence of SIV infection. We did not detect a consistent pattern of other proteins associated with activation, such as TIGIT, CTLA-4, HLA-DR, or ki67, on CD4+ or CD8+ T cells in PBMC from SIV+ animals compared to SIV-naïve animals (S2 Fig).

**Fig 6.**
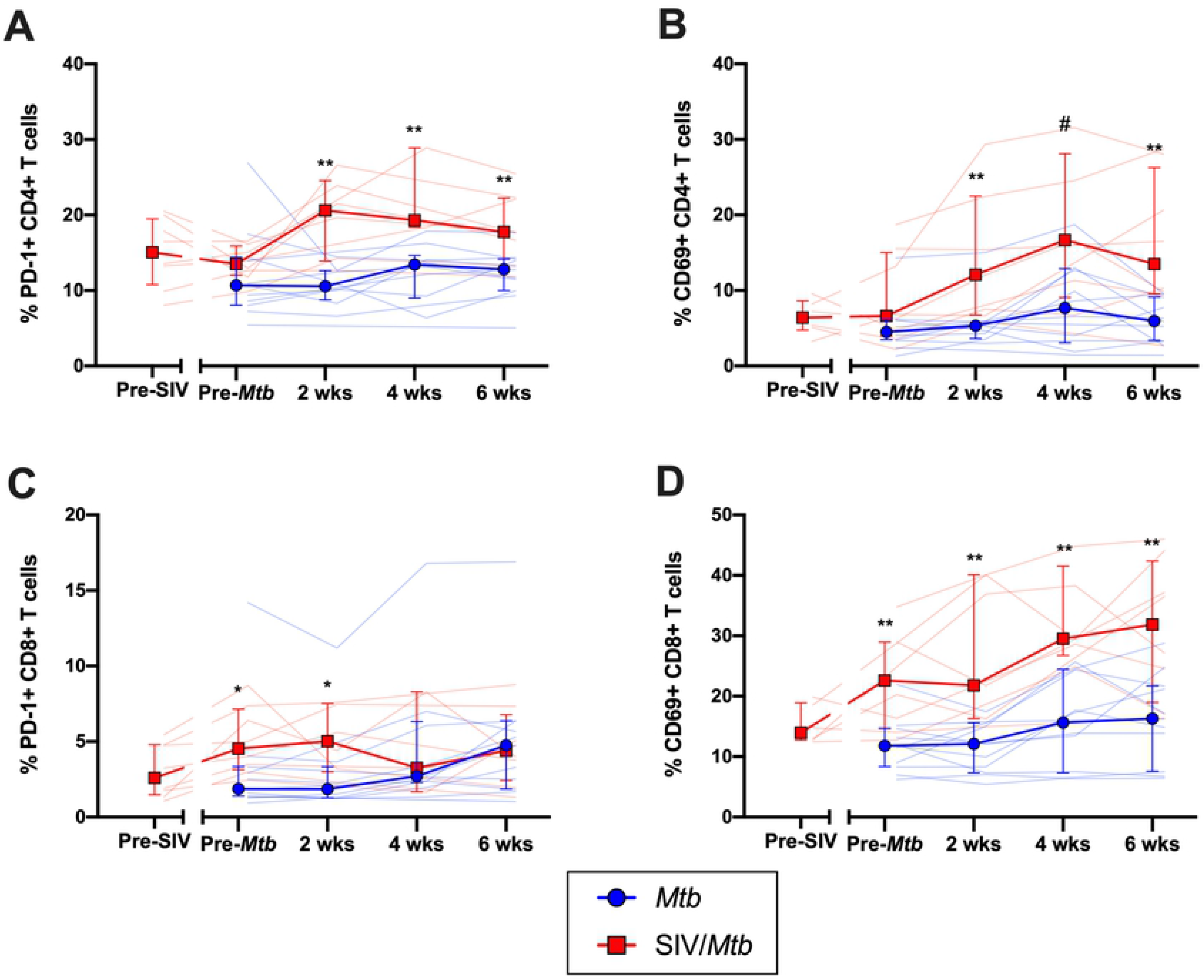
SIV+ animals have more circulating CD4+ and CD8+ T cells expressing PD-1 and CD69 following *Mtb* challenge. PBMC were collected and stained over the course of *Mtb* infection from *Mtb* (blue) and SIV/*Mtb* (red) animals. A.) PD-1 expression on CD4+ T cells. B.) CD69 expression on CD4+ T cells. C.) PD-1 on CD8+ T cells. D.) CD69 on CD8+ T cells. Population frequencies were determined by flow cytometry. Solid lines indicate median with interquartile range. Lighter lines indicate individual animals. Mann-Whitney tests were performed between SIV+ and SIV-naïve groups at each time point to determine significance; # 0.05 < p < 0.1, * p < 0.05, & ** p < 0.01.

We also compared T cell cytokine production in PBMC of SIV+ and SIV-naïve MCM following *Mtb* infection. We focused on two cytokines known to be important in anti-*Mtb* host immunity: IFN-γ and TNF. PBMC were stimulated with the immunogenic *Mtb* peptides ESAT-6/CFP-10 and subjected to intracellular staining and flow cytometry. We observed very little production of IFN-γ or TNF by CD4+ and CD8+ T cells in either SIV-naïve or SIV+ animals following *Mtb* infection (S3 Fig). This may reflect the very low frequency of T cells specific for these *Mtb* antigens present in blood, consistent with findings by others [42, 43]. Alternatively, this may be a consequence of using frozen PBMC which may reduce the number or function of antigen presenting cells required for inducing T cell responses [44].

### *Mtb*-specific T cells in airways of SIV+ animals produce less TNF following *Mtb* infection

*Mtb* transmission primarily occurs through inhalation, so immune responses in the airways is important to assess. We showed that CD4:CD8 T cell ratios in the airways were altered in SIV/*Mtb* co-infected MCM due to an influx of CD8+ T cells (Figs 2B and 4C). To test whether there was a difference in *Mtb*-specific T cells in the airways of SIV+ and SIV-naïve MCM following *Mtb* infection, BAL cells were stimulated *ex vivo* with pooled ESAT-6 and CFP-10 peptides and intracellular cytokine staining was performed (Fig 7A). These assays revealed no difference between SIV+ or SIV-naïve animals in the frequency of CD4+ or CD8+ T cells that produced IFN-γ alone (Fig 7B, C) or IFN-γ and TNF in combination (Fig 7D, E) after *Mtb* infection. However, production of TNF alone was decreased significantly 5 weeks post-*Mtb* in CD4+ and CD8+ T cells from SIV+ animals compared to SIV naïve animals (Fig 7F, G).

**Fig 7.**
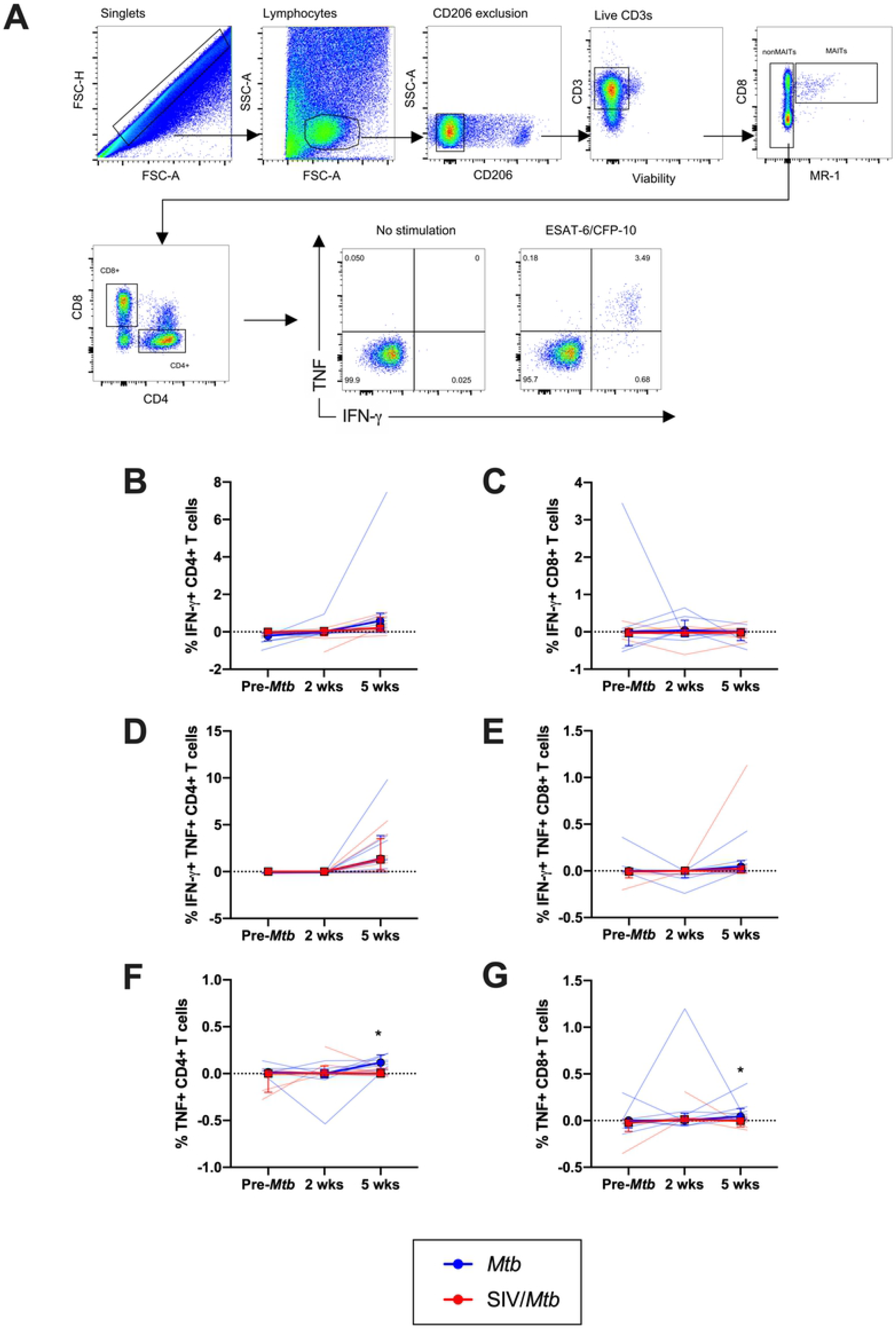
Decreased TNF production in response to *Mtb* antigen in BAL cells from SIV/*Mtb* animals. BAL were collected before *Mtb* and at 2 and 5 weeks post-*Mtb* challenge of *Mtb*-only (blue) and SIV/*Mtb* co-infected animals (red). Cells were stimulated for 5 hours with ESAT-6/CFP-10. A.) Example of BAL gating strategy. B.-G.) Frequency of CD4+ or CD8+ T cells producing IFN-γ and/or TNF. Frequencies were adjusted to background by subtracting from unstimulated controls. Solid lines indicate median with interquartile range. Lighter lines indicate individual animals. Mann-Whitney tests were used to determine statistical significance between SIV+ and SIV-naïve groups at indicated time points.

### SIV+ animals have increased frequencies of CD4+ and CD8+ T cells expressing PD-1 and TIGIT in lung tissue and granulomas

While BAL can be used to sample airway cells longitudinally, it cannot access cells within the lung parenchyma or within granulomas. We conducted comprehensive, PET/CT-guided necropsies 6 weeks after *Mtb* infection to harvest TB granulomas as well as random lung tissue samples to determine whether there were phenotypic and/or functional differences in these tissues between SIV+ and SIV-naïve animals.

Elevated immune activation in the lungs has been suggested previously to cause more severe TB disease in the setting of concurrent SIV infection [31]. Our hypothesis here was that the exacerbated TB disease that we showed previously in SIV+ MCM following *Mtb* co-infection [35] was due to SIV-associated immune activation present at the time of *Mtb* co-infection. Indeed, both lung tissue and granulomas harvested from SIV+ MCM 6 weeks after *Mtb* co-infection (red squares; Fig 8A, B) contained a higher frequency of CD4+ T cells expressing PD-1 than did those same tissues harvested from animals infected with *Mtb* alone (blue circles). There was no difference in the frequency of CD4+ T cells expressing PD-1 in lung tissue between SIV/*Mtb* co-infected animals (red squares; Fig 8A) and animals infected with SIV alone (black triangles) indicating that elevated PD-1 levels in SIV/*Mtb* animals was likely established prior to *Mtb* infection. Similarly, TIGIT expression was increased on CD4+ T cells in both lung tissue and granulomas of SIV/*Mtb* animals (red squares; Fig 8C, D) compared to those infected with *Mtb* alone (blue circles).

**Fig 8.**
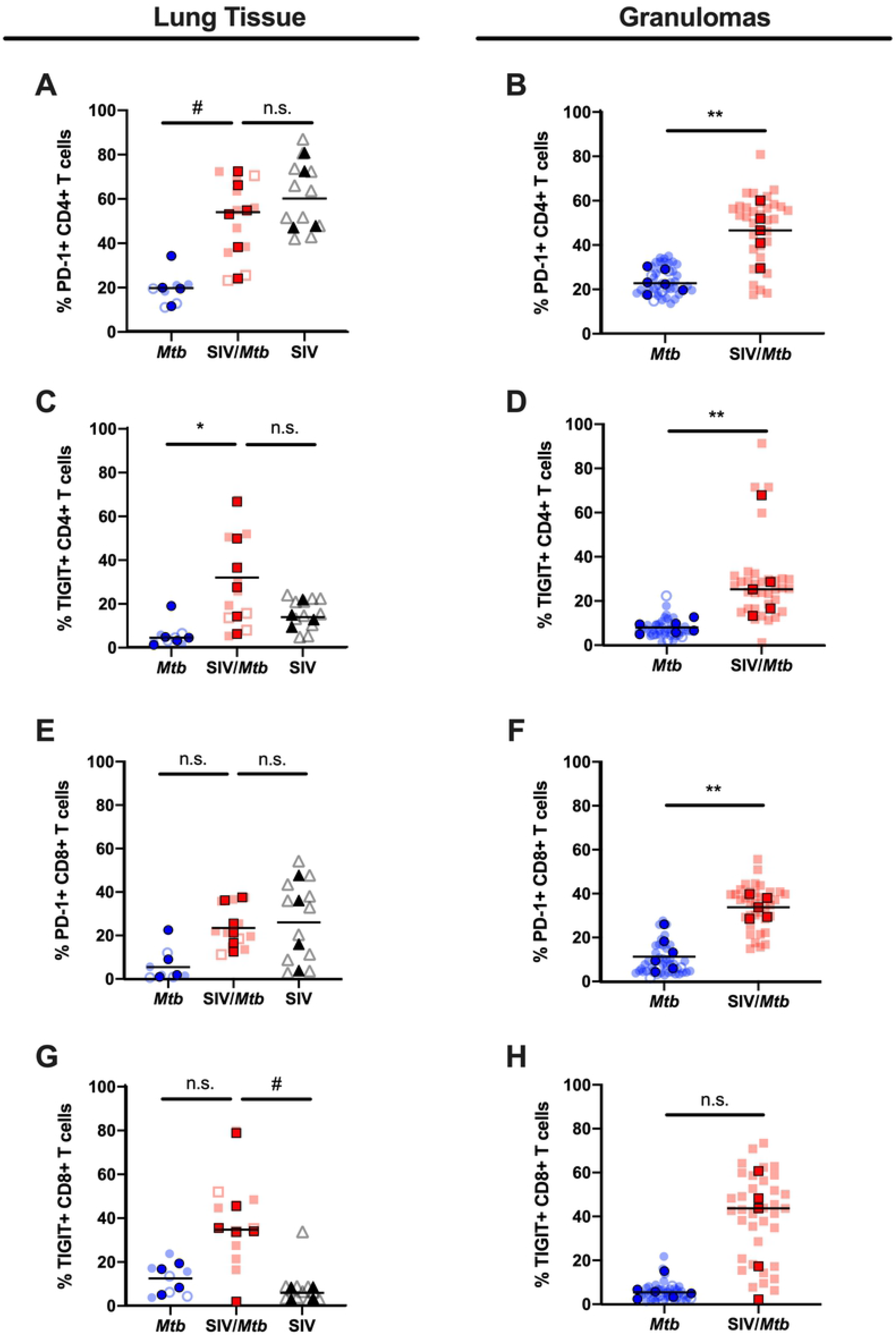
PD-1 and TIGIT expression is elevated in tissues of SIV+ animals. PD-1 and TIGIT expression on CD4+ and CD8+ T cells was measured by flow cytometry of tissues collected at necropsy for *Mtb*-only animals (blue) and SIV/*Mtb* co-infected animals (red). Darker symbols represent the median of each animal and the lighter symbols are individual samples. Closed symbols represent CFU+ tissue and open symbols are sterile (CFU-). Solid black line indicates median. For lung tissue (A., C., E., and G.), Kruskal-Wallis test was performed with a Dunn’s multiple comparison follow-up test comparing mean ranks to SIV/*Mtb*. For granulomas (B., D., F., and H.), Mann-Whitney tests were performed to determine statistical significance; # 0.05 < p < 0.1, * p < 0.05, & ** p < 0.01.

A higher frequency of CD8+ T cells expressing PD-1 was present in the granulomas of SIV/*Mtb* co-infected MCM (red squares; Fig 8F) compared to those infected with *Mtb* alone (blue circles). Conversely, there were no differences in the frequency of CD8+ T cells expressing PD-1 in lung tissue between animals infected with *Mtb* only (blue circles), SIV/*Mtb* (red squares), and SIV only animals (black triangles; Fig 8E). There was a trending increased frequency of CD8+ T cells expressing TIGIT in lung tissue of SIV/*Mtb* animals (Fig 8G, H).

### Granulomas from SIV+ animals contain lower frequencies of T cells producing TNF

The frequency of T cells producing the cytokines IFN-γ and TNF were measured in lung tissue and granulomas collected at necropsy 6 weeks after *Mtb* infection (Fig 9). IFN-γ production by CD4+ and CD8+ T cells differed little between SIV+ (red squares) and SIV-naïve (blue circles) animals in both lung tissue and granulomas (Fig 9A-D). However, there was a marked reduction in CD4+ and CD8+ T cells producing TNF in granulomas from SIV+ MCM (red squares) compared to SIV-naïve MCM (blue circles; CD4+, p = 0.0519; CD8+, p = 0.0519; Fig 9F, H).

**Fig 9.**
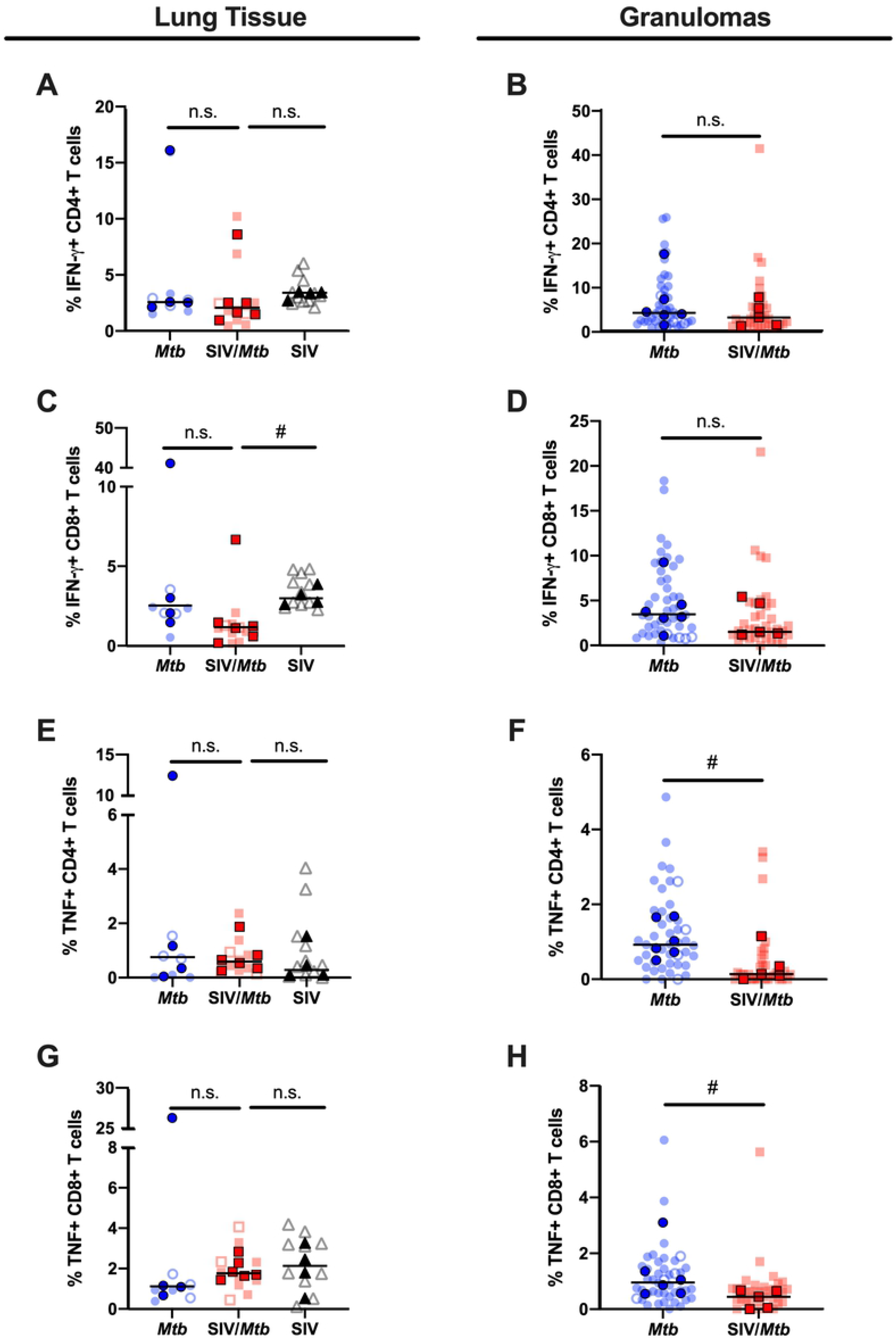
Less TNF production in granulomas of SIV+ animals. IFN-γ and TNF production of CD4+ and CD8+ T cells was measured by flow cytometry in tissues collected at necropsy for *Mtb*-only animals (blue) and SIV/*Mtb* co-infected animals (red). Darker symbols represent the median of each animal and the lighter symbols are individual samples. Closed symbols represent CFU+ tissue and open symbols are sterile (CFU-). Solid black line indicates median. For lung tissue (A., C., E., and G.), Kruskal-Wallis test was performed with a Dunn’s multiple comparison follow-up test comparing mean ranks to SIV/*Mtb*. For granulomas (B., D., F., and H.), Mann-Whitney tests were performed to determine statistical significance; # 0.05 < p < 0.1, * p < 0.05, & ** p < 0.01.

## Discussion

We previously demonstrated that, upon *Mtb* co-infection, animals with pre-existing SIV infection exhibit rapid TB disease progression, more severe TB disease, and earlier mortality than animals infected with *Mtb* alone [35]. We hypothesized that SIV caused immune dysregulation that led to exacerbated TB disease in SIV/*Mtb* co-infected animals. In this study, we sought to more precisely define that dysregulation by characterizing CD4+ and CD8+ T cell phenotypes and cytokine production in blood, airway, and tissues of SIV-infected macaques subsequently co-infected with *Mtb*, comparing to animals infected with *Mtb* alone.

We chose to conduct necropsies at 6 weeks post-*Mtb* infection since our initial study showed that TB progression, as measured by PET/CT imaging, began to diverge in SIV+ and SIV-naïve MCM between 4 and 8 weeks after *Mtb* infection [35]. Thus, the 6-week time point chosen here is the crucial time when adaptive immunity to *Mtb* is almost fully developed [14, 19] and the trajectories of TB disease are beginning to diverge in SIV+ and SIV-naïve animals, yet is prior to the appearance of advanced TB immunopathology (e.g. prolific necrosis, pneumonia, etc.) which we observed in our previous study [35] and which confounds careful immunologic characterization. Indeed, the SIV+ and SIV-naïve macaques necropsied at 6 weeks after *Mtb* infection exhibited similar TB pathology scores (Fig 1). They also had similar bacterial burdens at this time point (Fig 1), consistent with the work Lin, et al. who showed that mycobacterial killing capacity developed in granulomas sometime between 4 – 11 weeks post-infection [36]. We show here that, despite having comparable TB progression, SIV-naïve and SIV+ animals exhibited distinct immunologic differences following *Mtb* infection. Specifically, animals with pre-existing SIV had *i.* lower CD4:CD8 T cell ratios, *ii.* higher frequencies of CD4+ and CD8+ T cells expressing PD-1 or TIGIT at sites of *Mtb* infection, and *iii.* fewer T cells producing TNF in granulomas.

The initial immune response to *Mtb* infection is incredibly important in TB disease control [10]. Following *Mtb* infection, activated immune cells migrate to sites of infection to form granulomas, the hallmark of TB [10]. The granuloma is a complex, organized collection of cells consisting of macrophages, T and B lymphocytes, and neutrophils [10]. Of the lymphocyte population, T cells, both CD4+ and CD8+, are necessary to control *Mtb* infection [11, 12, 45–47]. HIV disrupts the balance between CD4+ and CD8+ T cells and some studies suggest that a reduction in the ratio of CD4:CD8 T cells is associated with poor TB disease outcome [38]. We found CD4:CD8 T cell ratios were lower across all tissue compartments measured in SIV/*Mtb* co-infected animals (Fig 2). A subset of animals chronically infected with SIV alone was necropsied to provide insight into the immunologic environment in tissues prior to *Mtb* infection. These animals also exhibited lower CD4:CD8 ratios in the blood and lung tissue, indicating that the decrease observed in SIV/*Mtb* animals was a consequence of their pre-existing SIV infection. In humans, HIV infection lowers CD4:CD8 T cell ratios in blood by a continuous loss of CD4+ T cells [48]. We observed a similar decline of circulating CD4+ T cells, without significant changes to CD8+ T cells, in SIV-infected animals, both before and after *Mtb* infection (Fig 3). Interestingly, in the airways, where immune cells first encounter inhaled *Mtb*, SIV/*Mtb* co-infected animals showed significant accumulation of CD8+ T cells (Fig 4). This is likely a consequence of SIV infection as SIV-specific CD8+ T cells have been shown to accumulate in airways following SIV infection [41].

The CD4:CD8 T cell ratios in both lung tissue and granulomas were lower in SIV/*Mtb* co-infected animals and was also noted in the lung tissue of SIV+ MCM, indicating that SIV infection alone drove the decline in the CD4:CD8 ratio (Fig 2). Looking more closely at the T cell populations in lung granulomas, we found an increase in CD8+ T cells present in granulomas in SIV+ animals compared to SIV-naïve animals (Fig 5B, right panel). The number of CD4+ T cells present in granulomas did not differ significantly between the two groups (Fig 5B, middle panel). Several groups have reported an influx of CD8+ T cells in airways of both HIV+ patients and SIV+ animals [30, 41, 49, 50]. One possible explanation for the accumulation of CD8+ T cells we observed in both BAL and granulomas of SIV/*Mtb* co-infected animals is that *Mtb* infection activates SIV-infected cells within the lungs and airways [51], increasing SIV replication, and driving an influx of the SIV-specific CD8+ T cells [41, 52]. While we did not examine whether certain functions by SIV-specific CD8+ T cells, such as cytokine secretion, impairs *Mtb* control within granulomas, future studies will investigate SIV replication in the airways and granulomas of *Mtb* co-infected MCM and assess SIV-specific CD8+ T cell responses to test this hypothesis.

We did not observe a difference in the frequencies of IFN-γ-producing CD4+ or CD8+ T cells within lung tissue or granulomas between SIV+ and SIV-naïve MCM at 6 weeks post-*Mtb* infection (Fig 9A-D). Nor did SIV+ and SIV-naïve MCM differ in the production of TNF by CD4+ and CD8+ T cells in lung tissue (Fig 9E, G). However, we did observe a decrease that trended towards significance in the frequency of CD4+ and CD8+ T cells producing TNF in granulomas from SIV+ animals compared to SIV-naïve animals. We also observed lower frequencies of TNF-producing CD4+ and CD8+ cells in the airways of SIV+ MCM in response to *Mtb*-specific stimuli at 5 weeks post-*Mtb* infection compared to SIV-naïve MCM (Fig 7F, G). Taken together, these data suggest that the diminution of TNF produced by CD4+ and CD8+ T cells as early as 5-6 weeks post-*Mtb* infection could explain, in part, the divergence in TB disease trajectory at later stages of *Mtb* infection. TNF is necessary for *Mtb* control [53–56]. TNF-deficient mice exhibit increased susceptibility to *Mtb* infection and rapidly succumb to *Mtb* infection [53–55]. In macaques, TNF neutralization resulted in exacerbation of active TB disease and reactivation of latent *Mtb* infection [56]. Moreover, patients using TNF-neutralizing agents are at a higher risk of acquiring TB or developing reactivated TB [57, 58]. Given the critical role of TNF in *Mtb* infection, future studies examining more carefully the role of this cytokine in SIV+ macaques upon *Mtb* co-infection are warranted.

Chronic immune activation is a feature of HIV and SIV infection that can be detrimental to host immunity [59–62]. In NHP species where SIV infection is non-pathogenic, immune activation resolves after several weeks following acute infection, while in species with pathogenic SIV infection, immune activation is maintained well into the chronic phase of SIV infection [62, 63]. Several studies have shown that unmitigated immune activation and cytokine production can lead to worsened TB disease [64, 65] and fails to enhance protection against *Mtb* [66–68]. In our SIV/*Mtb* animals, markers associated with an immune activation signature were elevated in blood and tissues when compared to animals infected with *Mtb* alone. In the blood from SIV+ MCM, we observed a higher frequency of both CD4+ and CD8+ T cells expressing markers associated with T cell activation (Fig 6). Interestingly, while peripheral CD4+ T cells were activated only after *Mtb* infection, CD8+ T cells were activated prior to *Mtb* infection in an SIV-dependent manner (Fig 6). In lung tissue as well as granulomas, there was a higher frequency of CD4+ T cells expressing PD-1 and TIGIT in both SIV/*Mtb* co-infected animals and animals infected with SIV only (Fig 8). These data indicate that CD4+ T cells in lungs of SIV+ animals are activated prior *Mtb* infection. We did not analyze whether cells expressing PD-1 or TIGIT produced less cytokine, but future SIV/*Mtb* co-infection studies will examine this as their expression of these receptors on T cells has been shown to have functional consequences on HIV/SIV- or *Mtb*-specific T cells [65, 69-77].

The immune events that occur during the early stages of *Mtb* infection are crucial in shaping the TB disease trajectory [10]. Our study found elevated expression of markers associated with immune activation on both CD4+ and CD8+ T cells in blood and tissues from SIV*+* animals. These markers remained elevated over the course of *Mtb* co-infection. Our data suggest that this SIV-induced immune activation may weaken the anti-mycobacterial function of granulomas, as reflected by a loss of TNF-producing CD4+ and CD8+ T cells just 6 weeks post-*Mtb* infection. These features may explain, in part, the increased susceptibility of PLHIV to TB.

## Materials and Methods

### Animal care and ethics statement

Adult (>4 years of age) Mauritian cynomolgus macaques (*Macaca fascicularis*; MCM) were obtained from Bioculture, Ltd. (Mauritius). MCM with at least one copy of the M1 MHC haplotype were selected for this study [78–80].

MCM that underwent *Mtb* infection (n=19) were handled and cared for at the University of Pittsburgh (U.Pitt.). MCM were housed in a BSL2+ animal facility during SIV infection and then moved into a BSL3+ facility within the U.Pitt. Regional Biocontainment Laboratory for *Mtb* infection. Animal protocols and procedures were approved by the U.Pitt. Institutional Animal Care and Use Committee (IACUC) which adheres to guidelines established in the Animal Welfare Act and the Guide for the Care and Use of Laboratory Animals, and the Weatherall report (8th Edition). The university is fully accredited by AAALAC (accreditation number 000496), and its OLAW animal welfare assurance number is D16-00118. The U.Pitt. IACUC reviewed and approved the IACUC study protocols 18032418 and 15035401, under Assurance Number A3187-01. The IACUC adheres to national guidelines established in the Animal Welfare Act (7 U.S.C. Sections 2131–2159) and the Guide for the Care and Use of Laboratory Animals (8th Edition) as mandated by the U.S. Public Health Service Policy. Macaques were housed at U.Pitt. in rooms with autonomously controlled temperature, humidity, and lighting. Animals were pair-housed in caging measuring 4.3 square feet per animal and spaced to allow visual and tactile contact with neighboring conspecifics. The macaques were fed twice daily with biscuits formulated for NHP, supplemented at least 4 days/week with fresh fruits, vegetables or other foraging mix. Animals had access to water *ad libitem*. An enhanced enrichment plan was designed and overseen by an NHP enrichment specialist. This plan has three components. First, species-specific behaviors are encouraged. All animals have access to toys and other manipulanda, some of which are filled with food treats (e.g. frozen fruit, peanut butter, etc.), and are rotated on a regular basis. Puzzle feeders, foraging boards, and cardboard tubes containing small food items also are placed in cages to stimulate foraging behaviors. Adjustable mirrors accessible to the animals stimulate interaction between cages. Second, routine interaction between humans and macaques are encouraged. These interactions occur daily and consist mainly of small food objects offered as enrichment and adhere to established safety protocols. Animal caretakers are encouraged to interact with the animals by talking or with facial expressions while performing tasks in the housing area. Routine procedures (e.g. feeding, cage cleaning, etc.) are done on a strict schedule to allow the animals to acclimate to a routine daily schedule. Third, all macaques are provided with a variety of visual and auditory stimulation. Housing areas contain either radios or TV/video equipment that play cartoons or other formats designed for children for at least 3 hours each day. The videos and radios are rotated between animal rooms so that the same enrichment is not played repetitively for the same group of animals. All animals are checked at least twice daily to assess appetite, attitude, activity level, hydration status, etc. Following SIV and/or *Mtb* infection, the animals are monitored closely for evidence of disease (e.g., anorexia, lethargy, tachypnea, dyspnea, coughing). Physical exams, including weights, are performed on a regular basis. Animals are sedated prior to all veterinary procedures (e.g. blood draws, etc.) using ketamine or other approved drugs. Regular PET/CT (Positron Emission Tomography/Computed Tomography) imaging is conducted on our macaques following *Mtb* infection and has proved very useful for monitoring disease progression. Our veterinary technicians monitor animals especially closely for any signs of pain or distress. If any are noted, appropriate supportive care (e.g. dietary supplementation, rehydration) and medications (e.g. analgesics) are given. Any animal considered to have advanced disease or intractable pain or distress from any cause is sedated with ketamine and then humanely euthanized using sodium pentobarbital (65 mg/kg, IV), consistent with the recommendations of the Panel on Euthanasia of the American Veterinary Medical Association (AVMA). Death is confirmed by lack of both heartbeat and pupillary responses by a trained veterinary professional. No animal on this study reached humane endpoint.

Four additional MCM were infected with SIV alone at the Wisconsin National Primate Research Center (WNPRC), where they were cared for in accordance with the regulations, guidelines, and recommendations outlined in the Animal Welfare Act, the Guide for the Care and Use of Laboratory Animals, and the Weatherall Report. The University of Wisconsin-Madison (UW-Madison), College of Letters and Science and Vice Chancellor for Research and Graduate Education Centers IACUC-approved the NHP research covered under IACUC protocol G005507. The UW-Madison Institutional Biosafety Committee approved this work under protocol B00000205. All macaques were housed in standard stainless-steel primate enclosures providing required floor space and fed using a nutritional plan based on recommendations published by the National Research Council. Macaques had visual and auditory contact with each other in the same room. Housing rooms were maintained at 65–75°F, 30–70% humidity, and on a 12:12 light-dark cycle (ON: 0600, OFF: 1800). Animals were fed twice daily a fixed formula, extruded dry diet with adequate carbohydrate, energy, fat, fiber, mineral, protein, and vitamin content (Harlan Teklad #2050, 20% protein Primate Diet, Madison, WI) supplemented with fruits, vegetables, and other edibles (e.g., nuts, cereals, seed mixtures, yogurt, peanut butter, popcorn, marshmallows, etc.) to provide variety to the diet and to inspire species-specific behaviors such as foraging. To further promote psychological well-being, animals were provided with food enrichment, structural enrichment, and/or manipulanda. Environmental enrichment objects were selected to minimize chances of pathogen transmission from one animal to another and from animals to care staff. While on study, all animals were evaluated by trained animal care staff at least twice daily for signs of pain, distress, and illness by observing appetite, stool quality, activity level, physical condition. Animals exhibiting abnormal presentation for any of these clinical parameters were provided appropriate care by attending veterinarians. Prior to all minor/brief experimental procedures, macaques were sedated with an intramuscular dose of ketamine (10 mg/kg) and monitored regularly until fully recovered from sedation. Per WNPRC standard operating procedure (SOP), all animals received environmental enhancement including constant visual, auditory, and olfactory contact with conspecifics, the provision of feeding devices which inspire foraging behavior, the provision and rotation of novel manipulanda (e.g., Kong toys, nylabones, etc.), and enclosure furniture (i.e., perches, shelves). At the end of the study, euthanasia was performed following WNPRC SOP as determined by the attending veterinarian and consistent with the recommendations of the Panel on Euthanasia of the AVMA. Following sedation with ketamine (at least 15 mg/kg body weight, IM), animals were administered at least 50 mg/kg IV or intracardiac sodium pentobarbital, or equivalent, as determined by a veterinarian. Death was defined by stoppage of the heart, as determined by a qualified and experienced individual.

### SIV and *Mtb* infection of MCM

At U. Pitt., animals in the SIV/*Mtb* co-infection group (*n* = 8) were infected intrarectally with 3,000 TCID50 of SIVmac239, as before [35]. After 6 months, the animals were co-infected with a low dose (3 - 12 CFU) of *Mtb* (Erdman strain) via bronchoscopic instillation, as described previously [81]. Animals in the SIV-naïve group (*n* =11) were infected similarly with *Mtb* alone. Six weeks post-*Mtb* infection animals were humanely euthanized as described above.

At UW-Madison, 4 MCM were infected intrarectally with 3,000 TCID_50_ SIVmac239. Six months after infection, animals were humanely euthanized as above.

### Clinical and microbiological monitoring

All animals were assessed twice daily for general health. Upon infection (or co-infection) with *Mtb*, animals were monitored closely for clinical signs of TB (coughing, weight loss, tachypnea, dyspnea, etc.). Monthly gastric aspirates and bronchoalveolar lavage (BAL) samples were tested for *Mtb* growth. Blood was drawn at regular intervals to measure erythrocyte sedimentation rate (ESR) and to serve as a source of peripheral blood mononuclear cells (PBMC) as well as plasma. In this study, no animal reached our humane endpoint criteria that included weight loss of >10%, prolonged cough, sustained increased respiratory rate or effort, and/or marked lethargy.

### Bacterial burden

To determine the number of live *Mtb* bacilli present in the lungs of each animal, a systematic approach [82] was used to plate tissue homogenates from every lung lesion, including both individual granulomas and complex pathologies such as consolidations, as well as from random pieces of grossly unaffected lung. Homogenates were plated on 7H11 medium agar (BD Difco), and *Mtb* CFU were enumerated after 21 days of incubation at 37°C and 5% CO_2_. Total lung bacterial load was calculated as described previously [82]. The total thoracic lymph node bacterial load was determined by harvesting all thoracic lymph nodes, regardless of whether pathology was grossly apparent, and plating as described above. The CFU from each sample were summed to yield total thoracic lymph node bacterial load. Summing the total lung and total thoracic lymph node CFU provided the total thoracic bacterial burden [82].

### PET/CT imaging

PET/CT scans were performed using a microPET Focus 220 preclinical PET scanner (Siemens Molecular Solutions) and a clinical eight-slice helical Ceretom CT scanner (NeuroLogica Corporation) as previously described [35, 82, 83], using 2-deoxy-2-(18F) fluorodeoxyglucose (FDG) as the PET probe. Each MCM was scanned at 4 weeks post-*Mtb* infection and again 1-3 days before necropsy. For each scan, individual granulomas were enumerated and lung inflammation was quantified by measuring total FDG activity [83].

### Sample collection

PBMC were isolated from whole blood drawn into Vacutainer tubes with EDTA (Becton Dickenson) at regular time points. Whole blood was centrifuged and plasma saved at −80°C for viral load analysis. Pellets were resuspended in phosphate-buffered saline (PBS; Lonza BioWhittaker) and layered over an equal volume of Ficoll (GE Healthcare). After centrifugation, the buffy coat was separated and washed with PBS. Contaminating red blood cells were lysed with Pharm Lyse (BD Biosciences). PBMC were resuspended in freezing medium containing 10% dimethyl sulfoxide and fetal bovine serum.

BAL was performed at time points throughout the course of SIV and/or *Mtb* infection by lavaging the airways up to four times with 10 mL of saline. The recovered sample was pelleted and BAL fluid was saved at −80°C. Pellets were resuspended in PBS, cells were enumerated using a hemocytometer and then immediately used for flow cytometry staining.

Necropsies were performed as previously described [35] at 6 weeks after *Mtb* infection. Within 3 days of necropsy, a final FDG PET/CT scan was performed to document disease progression and to provide a “roadmap” for collecting individual granulomas [82]. Animals were heavily sedated with ketamine, maximally bled, and humanely euthanized using sodium pentobarbital (Beuthanasia, Schering-Plough, Kenilworth, NJ). Granulomas matched to the final PET/CT images were harvested along with other TB pathologies (e.g., consolidations and pneumonia), thoracic and extrathoracic lymph nodes, and lung tissue (uninvolved or involved as determined *post facto* by *Mtb* culture), as well as portions of liver, spleen, mesenteric lymph nodes, ileum, and colon. Quantitative gross pathology scores were calculated to reflect overall TB disease burden for each animal [82]. Tissue samples were divided and a portion was fixed in 10% neutral buffered formalin for histopathology and the remainder was homogenized to a single-cell suspension as described previously [82]. Total cell counts of these single-cell suspensions were determined using a hemocytometer.

### Flow cytometry

To assess conventional CD4+ and CD8+ T cells, cells from PBMC, BAL, lung tissue, lymph nodes, and granulomas were stained as previously described [84]. Cell markers and antibodies are listed in Table 1. To assess frequencies of circulating T cell subsets, cryopreserved PBMC were used for staining. Corresponding whole blood samples were sent to the clinical hematology laboratory at the University of Pittsburgh Medical Center for complete blood counts (CBC). We used the flow cytometry data to convert the total lymphocyte numbers from the CBC to total CD4+ and CD8+ T cells/microliter of blood.

**Table 1.** Antibodies used in staining panels for flow cytometry.

| Marker | Clone | Fluorochrome(s) | Panel | Surface/Intracellular |
| --- | --- | --- | --- | --- |
| CD45 | D058-1283 | BV786 | Necropsy | Surface |
| CD3 | SP34-2 | BV650 | PBMC Stimulation, Necropsy | Surface |
| CD3 | SP34-2 | AF700 | PBMC Phenotype, BAL Phenotype | Surface |
| CD3 | SP34-2 | V500 | BAL Stimulation | Surface |
| CD4 | OKT4 | PE-Cy7 | PBMC Phenotype, BAL Stimulation, Necropsy | Surface |
| CD4 | L200 | BV510 | BAL Phenotype | Surface |
| CD4 | L200 | BV786 | PBMC Stimulation | Surface |
| CD8 | DK25 | PacBlue | BAL Phenotype, BAL Stimulation | Surface |
| CD8 | SK1 | BV510 | PBMC Stimulation | Surface |
| CD8 | RPA-T8 | BV711 | Necropsy | Surface |
| CD8 | RPA-T8 | BV786 | PBMC Phenotype | Surface |
| CD206 | 19.2 | PE-Cy5 | BAL Phenotype, BAL Stimulation, Necropsy | Surface |
| HLA-DR | G46-6 | BV650 | PBMC Stimulation | Surface |
| CD39 | eBioA1 (A1) | FITC | BAL Phenotype | Surface |
| CD25 | BC96 | BV605 | PBMC Stimulation | Surface |
| CD69 | TP1.55.3 | ECD | PBMC Stimulation | Surface |
| ki67 | B56 | AF647 | PBMC Phenotype, BAL Phenotype, Necropsy | Intracellular |
| TIGIT | MBSA43 | FITC | Necropsy | Surface |
| TIGIT | MBSA43 | PerCP-eFluor710 | PBMC Stimulation | Surface |
| PD1 | EH12.2H7 | BV605 | PBMC Stimulation, BAL Phenotype, Necropsy | Surface |
| CTLA-4 | 14D3 | PE-Cy7 | PBMC Stimulation | Surface |
| IFNγ | 4S.B3 | FITC | PBMC Stimulation, BAL Stimulation | Intracellular |
| IFNγ | 4S.B3 | BV510 | Necropsy | Intracellular |
| TNFα | Mab11 | AF700 | PBMC Stimulation, BAL Stimulation, Necropsy | Intracellular |
| IL-17A | BL168 | PE | Necropsy | Intracellular |
| CD107a | H4A3 | APC | BAL Stimulation | Surface |
| Live/Dead | -- | near-IR | PBMC Phenotype, PBMC Stimulation, BAL Phenotype, BAL Stimulation, Necropsy | Surface |

For analysis of BAL samples, freshly isolated cells were stained. Cells were resuspended in media (RPMI 1640 supplemented with 10% human albumin, 1% L-glutamine, and 1% HEPES) and divided for a phenotype flow panel and a stimulation flow panel (Table 1). The staining procedure for both panels were identical, except for the antibody cocktails used (Table 1). Approximately 1 × 10^6^ cells were stained per condition. Stimulated samples were stimulated with a mixture of peptides from the *Mtb* antigens ESAT-6/CFP-10 (1ug/mL pooled peptide each), or *Escherichia coli* (10 CFU) for stimulating MAIT cells [84], for 5 hours at 37°C, 5% CO_2_. Brefeldin A, monensin, and CD107a-APC were added 1 hour after the addition of stimuli. Following stimulation, cells were resuspended in 500 nM dasatinib (Thermo Fisher Scientific; Cat No. NC0897653) and stained with 0.25ug Mamu MR1 5-OP-RU or Ac-6-FP tetramer (PE-conjugated) for 1 hour (NIH Tetramer Core Facility, Atlanta, GA). Cells were washed with PBS + dasatinib and stained with Live/Dead™ near-IR (Thermo Fisher Scientific; Cat No. L10119) for 20 minutes in the dark at room temperature. Cells were washed with PBS + dasatinib and stained with the surface antibody cocktail (Table 1) for 20 minutes in the dark at room temperature. Cells were washed with FACS buffer (PBS + 1% FBS) supplemented with dasatinib and fixed overnight in 1% paraformaldehyde. For intracellular staining, cells were washed twice with FACS buffer and permeabilized with BD Cytofix/Cytoperm™ (BD; Cat No. 554714) for 10 minutes at room temperature, washed twice with BD Perm/Wash™ buffer, and stained with the intracellular cocktail (Table 1) for 20 minutes at room temperature.

To stain tissues obtained at necropsy, freshly isolated tissue homogenates were used. Approximately 1 × 10^6^ cells were stained when possible; granuloma homogenates often had < 1 × 10^6^ cells. Cells were resuspended in media as described above and were either left unstimulated or stimulated with phorbol 12,13-dibutyrate (PDBu) and ionomycin for 3 hours at 37°C, 5% CO_2_. The stimulators described above and brefeldin A were added simultaneously. Cells were then resuspended in 500 nM dasatinib and stained with 0.25ug Mamu MR1 5-OP-RU or Ac-6-FP tetramer (BV421-conjugated) for 1 hour. Within this incubation period, DRB*w501 tetramer (APC-conjugated) was added at 30 minutes (NIH Tetramer Core Facility, Atlanta, GA). Cells were subsequently washed with PBS + dasatinib and stained with Live/Dead™ near-IR for 20 minutes in the dark at room temperature. Cells were again washed with PBS + dasatinib and stained with surface antibody cocktail (Table 1) for 20 minutes in the dark at room temperature (Table 1). Cells were washed with FACS buffer supplemented with dasatinib and fixed overnight in 1% paraformaldehyde. For intracellular staining, cells were washed with FACS buffer and permeabilized with BD Cytofix/Cytoperm™ for 10 minutes at room temperature, washed twice with BD Perm/Wash™ buffer, and stained with intracellular cocktail (Table 1) for 20 min at room temperature.

Flow cytometry was performed on a BD LSR II (Becton Dickinson; Franklin Lakes, NJ), and the data were analyzed using FlowJo software for Macintosh (version 9.9.3 or version 10.1). Because cell numbers were often limited for granulomas, an event threshold was used to exclude samples with collected CD3 event counts ≤ 100. For calculating total number of cells in lung tissue and granulomas obtained at time of necropsy, total cell counts using a hemocytometer were multiplied by the specified population frequency relative to the lymphocyte gate. Lung tissue cell counts were reported as cells per gram of tissue, back-calculated from the sample weight at the time of collection. Granulomas with a cell count less than the limit of detection of the hemocytometer (5×10^4^ cells) were assigned a value of 4.5×10^4^ cells before correcting for the dilution factor as done previously [23].

### Statistical analysis

Due to their outbred nature, NHP studies are often hindered by substantial variation between individual animals. To avoid data bias by any one animal, medians of samples from individual animals were reported. The Shapiro-Wilk normality test was used to check for normal distribution of data. Pair-wise analysis of normally distributed data was performed using the unpaired *t* test. Nonnormally distributed data were analyzed with the Mann-Whitney test. For comparisons between multiple groups, the Kruskal-Wallis test was used. A Dunn’s multiple comparisons follow-up test was used to compare mean ranks of groups to the mean rank of SIV/*Mtb* co-infected animals. Statistical analysis for longitudinal data (PBMC and BAL) was performed in JMP 14 statistical software (version 14.0; SAS Institute). Statistical tests for all other data were performed in Prism (version 8.2.1; GraphPad). All tests were two-sided, and statistical significance was designated at a *P* value of < 0.05. *P* values between 0.05 and 0.10 were considered trending.

## Acknowledgements

We would like to thank our veterinary staff at the University of Pittsburgh and University of Wisconsin-Madison for their excellent animal care. We are grateful for the pathology expertise of Edwin Klein, DVM. We also thank the NIH Tetramer Core Facility for developing and providing the MR-1 5-OPRU and MR-1 6-FP tetramers.

## Supporting information

**S1 Fig. No difference in necropsy scores or CFU of individual compartments.** TB pathology was quantified in *Mtb*-infected animals at necropsy for individual compartments: lung (A.), thoracic lymph node (B.), and extrapulmonary (C.). *Mtb* bacterial load was determined for total lung (D.) and total thoracic lymph nodes (E.). The percent of granulomas that were CFU+ is shown in F. *Mtb*-only animals are depicted in blue; SIV/*Mtb* co-infected animals in red. Medians are shown and Mann-Whitney tests were used to determine statistical significance.

**S2 Fig. Activation/exhaustion marker expression on circulating CD4+ and CD8+ T cells over the course of *Mtb* infection.** PBMC were collected and stained over the course of *Mtb* infection from *Mtb*-only animals (blue) and from SIV/*Mtb* co-infected animals (red). Solid lines indicate median with interquartile range. Lighter lines indicate individual animals. Mann-Whitney tests were performed between SIV+ and SIV-naïve groups at each time point to determine significance; # 0.05 < p < 0.1, * p < 0.05, & ** p < 0.01.

**S3 Fig. No difference in cytokine response to ESAT-6/CFP-10 in CD4+ and CD8+ T cells over the course of *Mtb* infection.** PBMC were collected and frozen over the course of *Mtb* infection from *Mtb*-only (blue) and SIV/*Mtb* co-infected animals (red). Frozen PBMC were subsequently thawed, stimulated for 16 h with ESAT-6/CFP-10 and stained for flow cytometry. Frequencies were adjusted to background by subtracting values from unstimulated controls. Solid lines indicate median with interquartile range. Lighter lines indicate individual animals. Mann-Whitney tests were performed between SIV+ and SIV-naïve groups at each time point to determine significance.

